# Identification of Gly/NMDAR Antagonist from *Chromolaena odorata’s* Derived Phytoconstituents using Induced Fit Docking Approach

**DOI:** 10.1101/610006

**Authors:** Temitope Israel David, Olaposi Idowu. Omotuyi, Olalekan David Agboola, Dominic Chinedu Okonkwo, Niyi Samuel Adelakun

**Affiliations:** Centre for Biocomputing and Drug Development, Adekunle Ajasin University, Akungba Akoko, Ondo State, Nigeria; Department of Biochemistry, Adekunle Ajasin University, Akungba Akoko, Ondo State, Nigeria

**Keywords:** Antagonist, neurodegenerative, induced fit Docking, Lipinski rule of five

## Abstract

The ionotropic activation of N-methyl-D-aspartic acid (NMDA) plays a significant role in different type of neurodegenerative disease, as it is a tetramer with two Glycine binding subunit and two glutamate subunits. NMDA receptor can be inhibited by either blocking of the glycine site or glutamate site. Previously reported inhibitors of NMDA receptor focus on the inhibition of the glutamate subunit, which was reported to be associated with side effects such as ataxia, memory deficits, neurotoxicity. Therefore, different compounds with antagonistic effect are been explored on Gly/NMDA site. Glide XP docking was employed in screening phyto-constituent of *Chromolaena odorata* against Gly/NMDA receptor for hit compounds with antagonistic properties. The hit compounds were further subjected to Induced fit docking (IFD) and lipinski rule of five. The final selection was based on Rigid XP docking score using co-crystallized ligand as threshold docking score, interaction with receptor site residues, and IFD score. Ferulic acid, caffeic acid and scutellarein recorded binding affinity of −8.752Kcal/mol, 10.004 Kcal/mol and - 9.096 Kcal/mol respectively, which is higher than the binding affinity of co-crystallized ligand. Induced fit score obtained were −614.38, −614.03 and −616.31 for ferulic acid, caffeic acid and scutellarein respectively. The Adme properties of the phyto-constituent indicated that the compounds are drug-like in nature.

## Background

Activities of the central nervous system (CNS) are being controlled by excitatory and inhibitory amino acids which are known as neurotransmitters [1]. The balance between the excitatory and inhibitory are needed to maintain a healthy life whereas an imbalance in the two systems (excitatory and inhibitory) has been implicated in various mental disorders. Receptors for neurotransmitters are classified as ionotropic or metabotropic [1,2]. N-methyl-D-aspartic acid (NMDA) are excitatory ionotropic receptor that allows the passage of ions such as sodium, potassium and calcium. The influx of calcium is gated by NMDA receptor which is also known as the glutamate receptor existing in neurons. Glutamate being the most important excitatory neurotransmitter plays an important role in the pathophysiology of known degenerative disease of which they are responsible for the activation of ionotropic excitatory receptors such as NMDAR, AMPA and kainite receptors [2,3]. The activation of NMDA receptor by glutamate induces synaptic plasticity which is paramount for memory formation under normal circumstances, but can also be deleterious upon over stimulation of the receptor by abnormal release of glutamate which causes over stimulation of NMDAR in the post synaptic neurons [4,5]. This process has been implicated to be involved in excitotoxic neuronal damage in which there is excessive influx of calcium ion into cells leading to neuronal death [6,7].

NMDA receptor comprises of tetrameric assemblies of different subunit, in which the tetramer is formed from two subunit of glycine binding Glu1 and two subunit of glutamate binding Glu2. Its activity can be altered or blocked by interacting with glutamate agonist site, glycine agonist site, ion pores and allosteric sites on the amino acid terminal domain [3]. Previously developed NMDA receptor antagonist was based on interaction with the glutamate agonist site which has been proven to be effective but associated with various side effect profile such as neurotoxicity, ataxia and memory deficit [3,8]. Researches on co agonist Gly/NMDA receptor site showed that the site may be explored to circumvent side effects associated with the use of glutamate antagonist which is the main advantage of glycine site over glutamate co-agonist site [3]. Even with the role of Gly/NMDA receptor site in pathophysiology of neurodegenerative diseases, comprehensive structural study of component affecting fundamental activity is still not known. As a result of this research has been directed towards exploring different antagonist of Gly/NMDA receptor with little or no side effect profile.

Plants are made up of different chemical compounds known as phytochemicals, which confers different pharmacological properties on them. *C. odorata* has been reported to be made up of phytoconstituent such as the alkaloids, flavanols, polyphenol [9], some of which maybe of importance in inhibiting the Gly/NMDA binding site. Therefore, this study was designed to identify potential lead antagonist of Gly/NMDA from Chromolaena odorata phytoconstituent by induced fit docking. Induced fit docking was employed to allow flexibility of the active site amino acid residue around the lead compounds which can improve accuracy in the prediction of binding affinity of the antagonist at Gly/NMDA receptor site.

## METHODOLOGY

### Ligand preparation

Ligands of *Chromolaena odorata* were retrieved from reported journals, and their respective 2D structures of the ligands was downloaded from pubchem compound database in sdf format. The ligands were imported into maestro 11.5 interface where it was prepared using the ligand preparation tool(ligprep) [10]. Ligprep add missing hydrogen and converts the ligands into their respective 3D structure using OPLS3 force field at a pH of 7±2. On the ligprep interface, desalt and generate tautomers were selected and left to retain chirality, generating at most 32 per inputted ligands.

### Protein preparation and Site Generation

The co-crystallized 3-dimentional structure of Gly/NMDA receptor was acquired from protein data bank (PDB) with pdbid:1PBQ which was complexed with antagonist 5,7-dichlorokynurenic acid (DCKA) at 1.90Å [11]. The protein was selected with respected to low resolution and presence of an antagonist at its active site. The protein was imported into maestro 11.5 where it was prepared using the protein preparation wizard [12]. Protein preparation wizard is of importance in filling missing loops, side chains and addition of missing hydrogen atoms at a pH of 7±2. Het state was generated for the ligand which is followed by hydrogen bond assignment and refrain minimization.

Site generation defines the area around the binding site of ligand to receptor where interaction occurs. This was achieved using the receptor generation tool provided on maestro interface [13]. Receptor generation tool maps out the coordinate around the active site of the protein in x, y and z which encircles the bounded antagonist 5,7-dichlorokynurenic acid (DCKA). The grid generates a coordinate of X=5.61, Y=38.0 and Z= −17.16.

### Docking using Glide

Glide [14] was employed in analyzing the binding affinity between the library of compounds and the active site. In this case, the active site is held rigidly around the docked ligands with the ligands assuming different pose at the active site. The co-crystallized ligand and prepared ligands were selected from the project table and docked initially using the standard precision algorithm (SP) with ligand sampling selected as flexible, followed by extra precision algorithm (XP) with ligand sampling at none refined only. The binding energy of library of ligands of *Chromolaena odorata* was compared with that of co-crystallized ligand (5,7-dichlorokynurenic acid) with the binding energy of co-crystalized ligand as cutoff binding score.

### Induced Fit Docking (IFD)

Induced fit docking protocol was performed on ligands using the induced fit tool in maestro 11.5 [15]. IFD protocol allows the flexibility of both the receptor of the protein and ligand, which has been reported to be robust and very accurate in predicting binding affinity between ligand and active site of a protein [16]. IFD employs the use of glide and prime for ligand docking and protein refinement respectively. The active site was centered on co-ordinate A: 1001 to include the amino acid residues around 5, 7-dichlorokynurenic acid of 1PBQ. The inputted ligand was set to sample ring conformation at an energy window of 2.5kcal/mol. The initial docking protocol was set to employ van dar aal scalling of 0.5 for both the ligand and receptor and to generate maximum of 20 poses per ligand. Induced fit protein-ligand complexes was generated using prime and further subjected to side chain and backbone refinement. Extra precision algorithm (XP) was employed in redocking of the ligand with the low energy refined strictures generated by prime. XP glide score, and IFD score was computed for each of the protein-ligand complexes which accounts for the protein-ligand interaction energy and the total energy of the system. Complex with more negative induced fit score has a more favorable binding and the interaction was viewed with 2d interaction diagram tool in maestro 11.5

### Validation of Docking Protocol

The docking procedure was validated by calculating the RMSD value of pose obtained before and after docking procedure of 5,7-dichlorokynurenic acid as shown in Figure 2.

### Pharmacokinetic parameters

The pharmacokinetics properties of the hit compounds were evaluated using the qikprop program embedded in maestro 11.5 [17]. This is crucial in eliminating toxic and compounds with low probability of reaching the target protein.

## Result and Discussion

Molecular docking enables the screening of large database of compounds against target protein to identify possible hit compounds [18]. This procedure is based on the ability of the compounds to interact with amino acid residues, which is based on the protein conformation, and the assumed pose of the ligand [18]. Library of phytochemicals generated was able to interact perfectly with binding affinities lesser than the co-crystallized ligand showing their antagonistic effect on Gly/NMDA receptor. The extra precision docked score obtained is because the receptor of the protein is held rigid around the ligands, which might give a result that might not be reliable if the ligands is able to induce a conformational change at the receptor site. Therefore, induced fit docking was employed on the hit compounds to give room for flexibility of the receptor using prime program. The hit induced a significant change on the receptor giving room for a better docking score and interaction with the amino acid residues when compared to result obtained via rigid docking using glide. The sum total of these interactions is used to compute an induced fit score for the bound ligand. In this case, ferulic acid and caffeic acid recorded an induced fit score of −614.38 and −614.03 respectively when compared to the co-crystallized ligand which recorded an induced fit score of −611.13. Admet/tox screening was also recorded for the ligands which indicate their effectiveness as Gly/NMDA antagonist. The docked score, induced fit score and admet/tox screening scores obtained from interaction of hit compounds from library of phytochemicals of *Chromolaena odorata* are shown in Table 1.

**Table 1:**
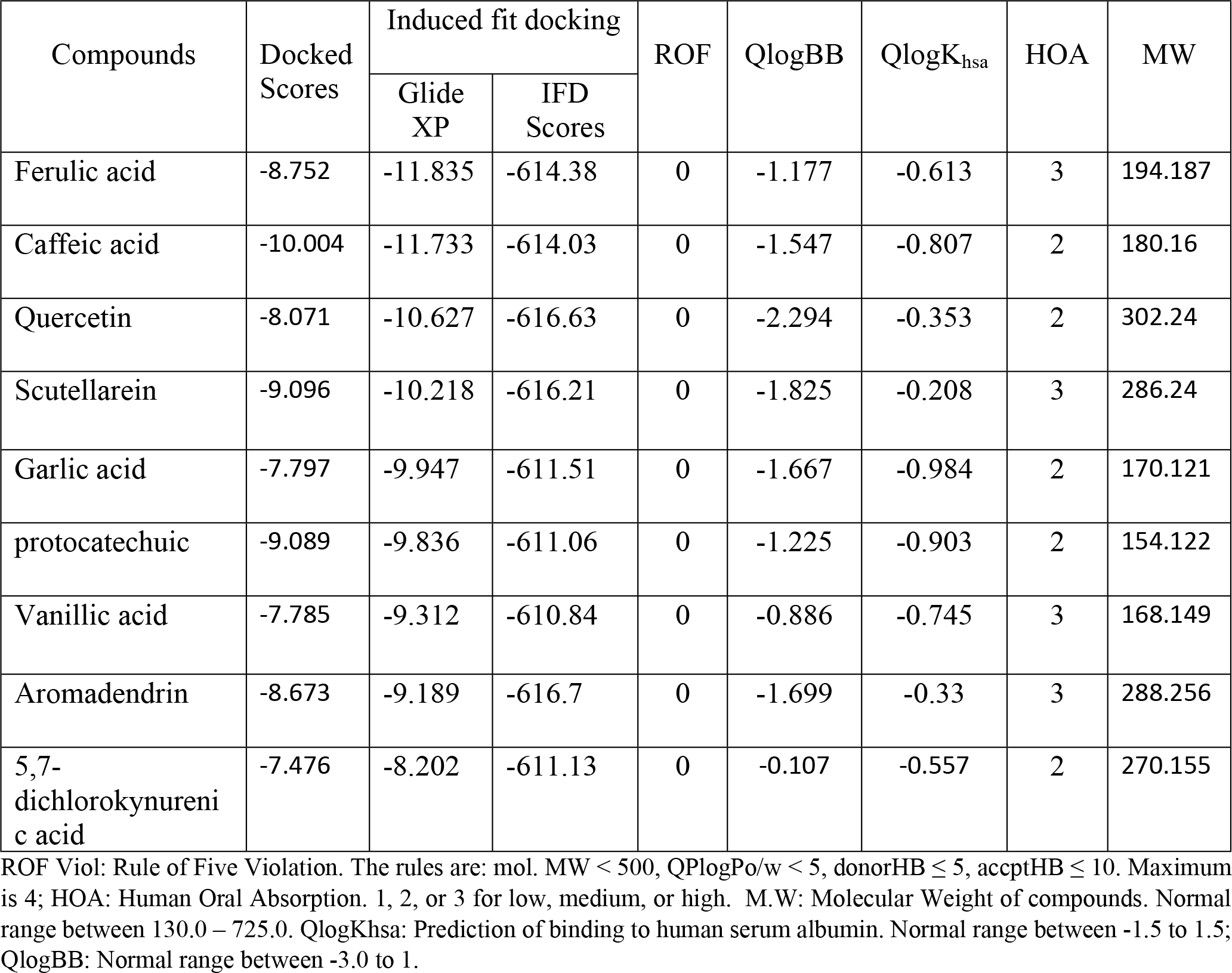
Showing the IFD, ADME screening and XP rigid docking of *Chromolaena odorata* to Gly/NMDA receptor

### Interaction profiles

Ligands at the active site of proteins are stabilized by interactions with the amino acid residues in which this interaction contributes significantly to the binding energy of the ligands. The pose of the ligands also contributes to the binding attribute of the ligand. Induced fit docking enables the best predicting ability of ligands using the combination of protein flexibility around the ligand and glide docking, and redocking of the ligand to obtain best interaction and orientation of ligand at the receptor site [18]. The 2d interaction of hit ligand at the active site of Gly/NMDA receptor is represented with 2D stick on figure 1. The ligands exhibit similar interaction with the co-crystallized ligand with the formation of hydrogen bond, hydrophobic interactions and pi-pi interactions which is similar to those recorded by Ugale and Baari (2016) [3]. Ugale and Bari recorded antagonistic interaction of NMDA inhibitors with amino acid residues Arg131, Pro124 and Thr124 [3]. This interactions was validated by Sharma and Gupta (2012) [1] which recorded similar interactions and some additional interactions with Thr126 an Trp223. Ferulic acid and caffeic acid assumes similar orientation, being enclosed by hydrophobic site residues. Ferulic acid, Quercetin, vanillic acid and caffeic acid form hydrogen bond with hydroxyl group attached to the phenyl ring with negatively charged residue Arg244 and Trp223. The carboxyl group of the acidic ligand (ferulic acid, caffeic acid, garlic acid, vanillic acid and protocatechuic) is buried deep within the hydrophobic pocket forming hydrogen bond and a salt bridge with positively charge residue Arg131, and hydrogen bond acceptor from polar side Thr124. The phenylic ring of garlic acid, vanillic acid, protocatechuic and scutellarein forms pi-pi stacking with hydrophobic side residue Phe92 that might be crucial for the stability of the ligands at the binding site. Quercetin, scutellarein and aromadenderin shares a common structure with two phenyl rings joined by heterocyclic ring, forming hydrogen bond using their carbonyl group attached to their heterocyclic with positively charged Arg131 and Thr126 in case of scutellarein. Hydrogen bond is formed between polar Gln13 with hydroxylphenyl group of Gallic acid and aromandedrin. Quercetin and aromandedrin recorded similar orientation at the active site of Gly/NMDA forming hydrogen bond interaction with Pro124 and Thr126 using the hydroxyl group attached to their heteroxylxlic ring.

**Figure 1:**
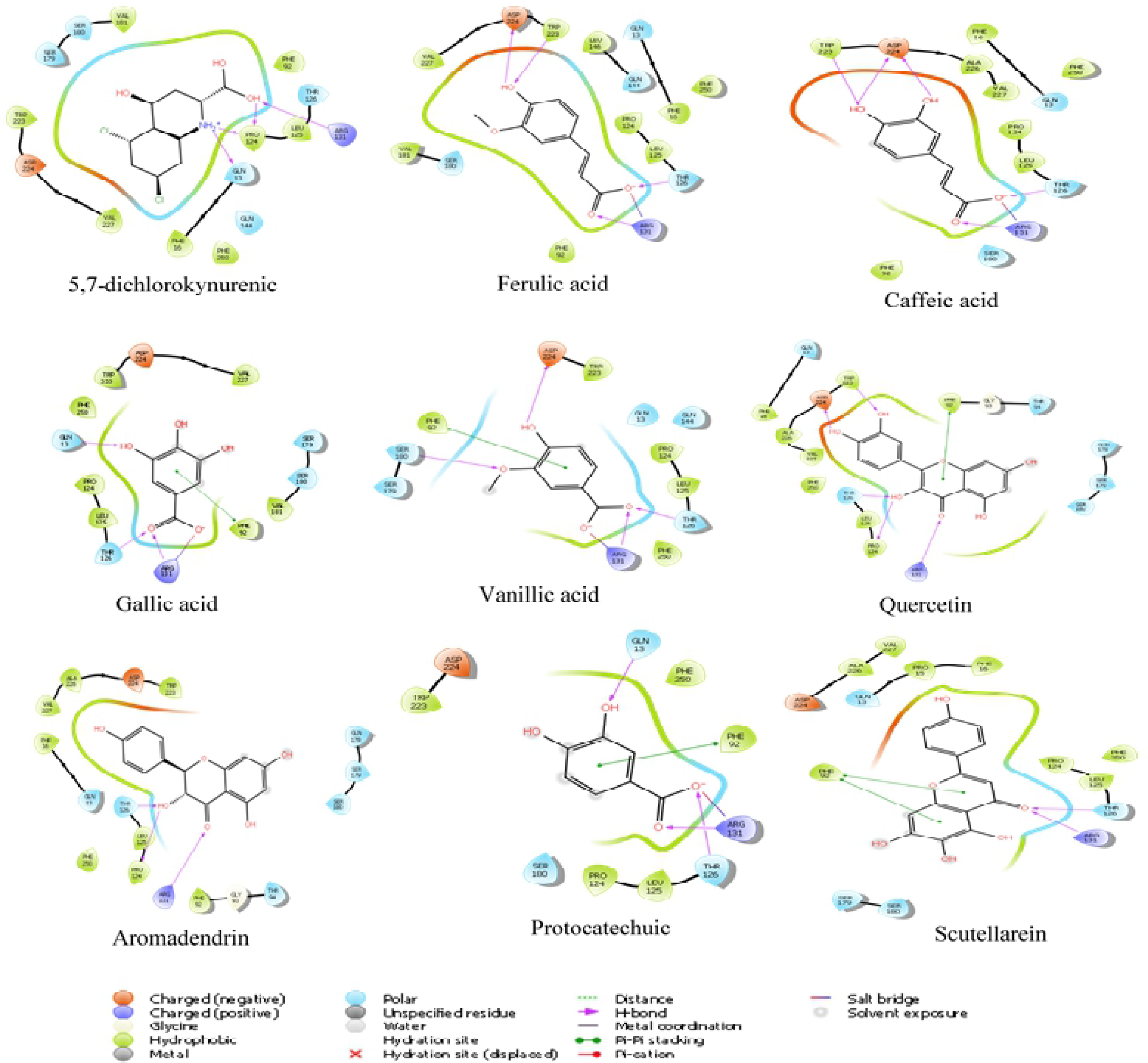
2D interaction profile of the phyto-ligands with specific amino acid residues of NMDA receptor

**Figure 2:**
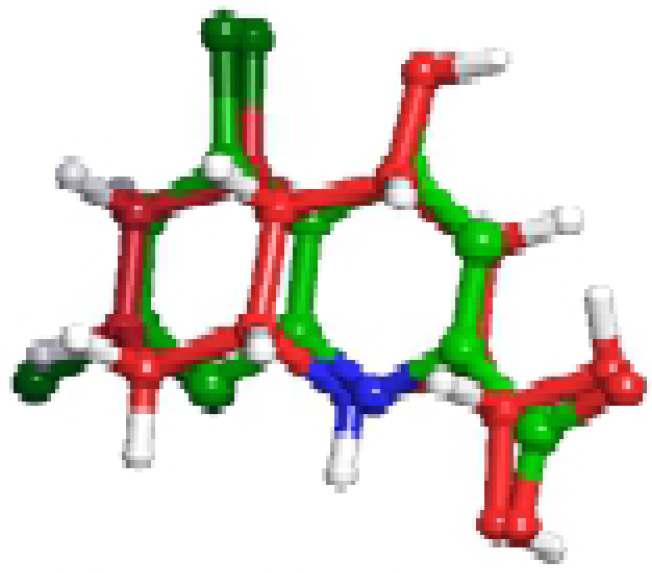
Showing the validation of the docking protocol employed with co-crystallized ligand (green) before docking and after glide XP docking (Red). The co-crystallized ligand overlaps almost perfectly with an RMSD value of 0.8298 indicating accuracy in the docking protocol employed.

## ADME/TOX

Evaluation of the pharmacokinetic property of ligands is useful in screening a large database of compounds for hit compounds. The tool is also useful in eliminating toxic ligands and evaluating the drug likeness properties of a compound. The hit compounds were evaluated using parameters such as the absorption, distribution, metabolism and elimination (ADME) [19]. These parameters is based on the lipinski rule of five (ROF) which enlist important criteria needed before a compound to be considered to be drug like. From our adme study, the hit compounds violated none of the lipinski rules, which qualifies them to possess drug like properties. Some important parameters were also taken into consideration such as the human oral absorption, binding of the ligands to human serum albumin and blood brain barrier, with most of the ligands recording a high human oral absorption. In addition, the ligands fall between the normal ranges of their binding with human serum albumin which shows their effectiveness in binding to target receptor. The activity of these ligands on the CNS is also crucial, and it was deduced that the ligands recorded a low activity of the CNS which falls within normal range of −3.0-1.0.

### Conclusion

It is of paramount importance to explore natural sources for compounds with druglike properties, which confers less side effect when compared to synthetic drugs. This study has shown the ability of phytochemicals from *Chromolaena odorata* to act a potent antagonist of Gly/NMDA receptor, which can be utilized in treatment of neurodegenerative diseases. Therefore, it is required that further research be conducted on the ability of these compounds to be a potent antagonist of Gly/NMDA receptor.

## Acknowledgements

I want to acknowledge the contribution of center for bio-computing and drug development (CBDD) Adekunle Ajasin University for their mentorship towards the writing of this manuscript

